# Effects of Short-time Exposure to Atrazine on miRNA Expression Profiles in the Gonad of Common Carp (*Cyprinus carpio*)

**DOI:** 10.1101/345371

**Authors:** Fang Wang, Qian-wen Yang, Wen-Jie Zhao, Qi-Yan Du, Zhong-Jie Chang

## Abstract

MicroRNAs (miRNAs) are endogenous small non-coding RNAs that negatively regulate gene expression by targeting specific mRNAs; they are involved in the modulation of important mRNA networks involved in toxicity. Atrazine is a known endocrine-disrupting chemical, whose molecular mechanisms are unknown. In this study, common carp (*Cyprinus carpio*) gonads at two key developmental stages were exposed to 0.428 ppb atrazine for 24 h *in vitro*. MiRNA expression profiles were analysed to identify miRNAs related to gonad development and to reveal the atrazine mechanisms interfering with gonad differentiation. Atrazine exposure caused significant alteration of multiple miRNAs. Compared with the juvenile ovary, more miRNAs were down-regulated in juvenile testis, some of these down-regulated miRNAs target the steroid hormone biosynthesis pathway related-genes. Predicted target genes of differently-expressed miRNAs after exposure to atrazine were involved in many reproductive biology signalling pathways. We suggest that these target genes may have important roles in atrazine-induced reproductive toxicity by altering miRNAs expression. Our results also indicate that atrazine can up-regulate aromatase expression through miRNAs, which supports the hypothesis that atrazine has endocrine-disrupting activity by altering the expression of genes of the Hypothalamus-Pituitary-Gonad axis through its corresponding miRNAs. This study tells us the following conclusions: 1. Atrazine exposure results in significant alterations of miRNAs whose predicted target genes are associated with reproductive processes. 2. In the primordial gonad, atrazine promoted the expression of early gonad-determining genes by decreasing specific miRNAs. 3. In the juvenile gonad, atrazine promoted the biosynthesis of steroid hormones.

## INTRODUCTION

Sex determination in fish is significantly influenced by environmental factors, such as temperature, pH, exogenous hormones, and pollutants (Devlin et al., 2002). Pollutants, such as pesticides, are potential endocrine disruptors, which even at very low levels are sufficient to cause developmental and reproductive alterations in numerous species (Colborn et al., 1993; Corcoran et al., 2010).

With the development of agriculture, herbicides are increasingly used to reduce soil erosion, to avoid the manual removal of weeds, and to increase crop production rates (Gianessi & Sankula, 2003). However, the use of pesticides leads to serious harm to living organisms. Atrazine (2-chloro-4-ethylamino-6-isopropylamino-1,3,5-triazine) is a pre-emergent herbicide used on a variety of agricultural crops including corn, sorghum grass, sugar cane, and wheat (Barr et al., 2007; Eldridge et al., 2008; Solomon et al., 2008). Atrazine is probably the most widely used herbicide in the world frequently contaminating potable water supplies (U.S. Environmental Protection Agency 1994). Atrazine is a suspected endocrine-disrupting chemical that alters male reproductive tissues, when animals are exposed during development.

Various studies indicate atrazine adversely impacts the neuroendocrine and reproductive systems, and that it may be a potential carcinogen (Cooper et al., 2007; Cragin et al., 2011; Hayes et al., 2010; Freeman et al., 2011). Currently the epigenetic, genetic, and cellular mechanisms altered by atrazine exposure are under investigation (Kucka et al., 2012; Karmaus and Zacharewski, 2015; Pogrmic et al., 2009, Pogrmic-Majkic et al., 2010, 2014; Wirbisky et al., 2016a,b). Tevera-Mendoza et al. (2002) showed that atrazine exposure, for as little as 48 h at 21 ppb, resulted in severe gonad dysgenesis in African clawed frogs (*Xenopus laevis*). Moreover, atrazine induced hermaphroditism at concentrations of only 0.1 ppb (Hayes et al. 2002). In fish, atrazine can result in complete feminization of males, as illustrated by skewed sex ratios in zebrafish (*Danio rerio*), which have no distinguishable sex chromosomes (Suzawa et al., 2008).

In zebrafish, atrazine exposure during embryonic development alters MicroRNAs (miRNAs) associated with angiogenesis, cancer, and neurodevelopment (Sara et al., 2016). Numerous studies have shown that atrazine has adverse effects on the neuroendocrine system, primarily affecting the hypothalamus–pituitary–gonad (HPG) axis. Atrazine decreases gonadotropin-releasing hormone release, the pre-ovulatory surge of luteinizing hormone, follicle stimulating hormone, and prolactin (Cooper et al., 2000; Foradori et al., 2009, 2013; Weber et al., 2013; Wirbisky et al., 2016a). However, the mechanism of action of atrazine is not well-understood.

MiRNAs are single-stranded, highly conserved, non-coding RNA molecules of 19–24 nucleotides (nt), which regulate gene expression at the post-transcriptional level, by targeting specific sites in the 3’ untranslated region of mRNAs (Bartel, 2004; He et al., 2004; Krol et al., 2010). miRNAs play important roles in controlling multiple biological processes, such as embryonic development, cell cycle control, apoptosis, cell proliferation and differentiation, and immune and stress responses in various organs (Brennecke et al., 2003; Hwang et al., 2006; Pedersen et al., 2007; Ro et al., 2007; Xu et al., 2003). In the last few years, miRNAs have been reported to play an important role in the response to toxicant exposure and in the process of toxicant-induced tumorigenesis (Jardim et al., 2009; Rager et al., 2011; Zhang and Pan, 2009).

As a new tool for risk assessment, miRNAs can provide indications on the toxicology mechanisms associated with environmental factors and with disease. MiRNAs are also novel biomarkers of the diseases related to environmental factors (Li et al., 2014). Recently, an increasing number of studies have shown that miRNAs can functionally interact with a variety of environmental factors including drugs, viruses, radiation, and environmental chemicals (e.g., formaldehyde, PAHs, and bisphenol A) (Izzotti and Pulliero, 2014; Qiu et al., 2012; Ray et al., 2014). Knowledge on the role miRNAs in toxicological responses is increasing, but is still limited.

The common carp, *Cyprinus carpio*, is one of the most important cyprinid species, accounting for 10% of the global freshwater aquaculture production (Xu et al., 2014). Genomic studies of common carp have recently made extensive progress. Common carp transcriptome was deep sequenced by Ji et al. (2012) and Jiang et al. (2016), who identified changes at the transcriptomic level in common carp spleen after 24 h of experimental infection with *Aeromonas hydrophila*. A large number of gene associated single-nucleotide polymorphisms (SNPs) were identified in four strains of common carp using nextgeneration sequencing (Xu et al., 2014). miRNAs and miRNA-related SNPs were also identified. MiRNA-related SNPs affect biogenesis and regulation in the common carp (Zhu et al., 2012).

Yellow River carp (common carp from the Yellow River) is famous in China for its tender, tasty, and nutritional meat. Females grow faster than males, which makes the mechanism of sex differentiation and development an intriguing topic in this commercially important species (Gui et al., 2012; Mei and Gui, 2015). In our previous study, we profiled miRNAs from five different developmental stages of Yellow River carp, in order to identify differentially-expressed and novel miRNAs that may play regulatory roles in ovary differentiation (Wang et al., 2017). Our previous study showed that there is a dynamic shift in gene expression during gonad differentiation and development. (Jia et al., 2017). Environmental factors can affect miRNAs in fish, and even play a decisive role in some species.

Several studies have shown that in zebrafish and humans atrazine exposure alters miRNAs associated with angiogenesis, cancer, and neurological development (Wirbisky et al., 2016). However, few studies have investigated the role of miRNAs in toxicological responses during sex differentiation and development in teleost fish.

In this study, we looked for correlations of miRNA and mRNA expressions during sex differentiation and development of carp, following atrazine exposure. The gonad development of carp has several critical periods, including primordial gonad and juvenile gonad. It would be valuable to understand the gene expression changes and the roles of miRNAs during the key stages of gonad development of carp, when they are exposed to atrazine. Therefore we aimed to investigate the effect of atrazine exposure on the global expression profile of miRNAs in the two key stages of gonad development by deep sequencing. We also predicted target genes that would affect gonad development. Our results would help us to better understand the molecular mechanisms of atrazine toxicity on gonad development, and to reveal the roles of miRNA–mRNA interactions in toxicological mechanisms, and the important impact on sex differentiation and gonad development of common carp.

## MATERIALS AND METHODS

### Chemicals

Atrazine (purity > 98%) was purchased from Beijing Dezhong-Venture Pharmaceutical Technology Development Co., Ltd. (Beijing, China). As atrazine has low solubility in water, the stock solutions and dilutions were prepared in acetone (Fisher Scientific, USA) and stored at 4 °C.

### Fish Samples

All investigations in this study were performed according to the Animal Experimental Guidelines of the Ethical Committee of the University of China. The Yellow River carp used in this study were obtained from the aquaculture facilities of Henan Normal University and maintained at the genetics laboratory (Henan normal university, Xinxiang Henan province, China) in flow-through water tanks with a constant temperature of 25 ± 1 °C. The test samples included gonads from two different developmental stages. Samples of primordial gonads were collected from larvae at 45 days post-hatching, based on the results of our previous studies (Wang et.al, 2017). The original reproductive gland was dissected under a microscope, and samples from 50 fish were mixed after confirmation by histological section. Samples of juvenile gonad were collected from 30 fish 80 days post-hatching. Stage II ovaries and testis were confirmed with histological sections.

### Atrazine Exposure

Samples of two different stages including primordial gonad and juvenile gonad (ovary and testis) were cultured at 28 °C in a humidified 10% CO_2_ atmosphere in Dulbecco’s modified eagle medium supplemented with 10% foetal bovine serum (Gibco, Life Technologies) (Pombinho et al., 2004; Daniel et al., 2014). Culture medium was renewed every two days. For atrazine exposure experiments, cells were seeded in 24-well plates and allowed to proliferate for 48 h. Then samples were treated with 0.428 ppb of atrazine for 24 h. Three replicates were set for each treatment, as well as for the unexposed control. Samples were collected at 8 h and 24 h post-treatment, and were immediately frozen in liquid nitrogen and stored at −80 °C for further use.

### RNA Isolation

Total RNA was extracted from each sample separately using TRIzol reagent (Invitrogen, Carlsbad, CA, USA) following the manufacturer’s protocol. The quantity and purity of total RNA were checked using the Agilent 2100 Bioanalyzer system (Santa Clara, CA, USA) and by denaturing gel electrophoresis. The samples were then stored at −80 °C.

### Small-RNA Library Construction and Sequencing

We generated small-RNA libraries from the nine samples from Yellow River carp: primordial gonad control (PG-CK), primordial gonad exposed to atrazine for 8 h (PG-A8h), primordial gonad exposed to atrazine for 24 h (PG-A24h), juvenile ovary control (IIC-CK), juvenile ovary exposed to atrazine for 8 h (IIC-A8h), juvenile ovary exposed to atrazine for 24 h (IIC-A24h), juvenile testis control (IIX-CK), juvenile testis exposed to atrazine for 8 h (IIX-A8h), juvenile testis exposed to atrazine for 24 h (IIX-A24h). Small-RNA libraries were generated using the mirVanaTM mircoRNA Isolation Kit (Ambion, USA), according to the manufacturer’s instructions. Small-RNA libraries were prepared from three biological replicates for each sample.

Total RNA was ligated with 3′ and 5′ RNA adaptors. Fragments with adaptors on both ends were enriched by PCR after reverse transcription, as described previously (Wang et al., 2017). The resulting cDNAs were purified and enriched with 6% denaturing polyacrylamide gel electrophoresis to isolate the fractions of the expected size and to eliminate unincorporated primers, primer di mer products, and dimerized adaptors (Wang et al., 2017). Finally, the nine resulting RNA libraries were sequenced using an Illumina/Solexa Genome Analyzer, at Guangzhou Genedenovo Biotech Company (Guangzhou, China).

### Sequencing Data Analysis

As we described previously (Wang et Al., 2017), the raw sequence data were filtered to remove low quality reads and adaptor sequences. After adaptor trimming, reads of 16–35 nt in length were kept for further bioinformatic analysis. The remaining reads were mapped to the *C. carpio* genome with a tolerance of zero mismatches in the seed sequence using Bowtie (version 1.1.0). Sequences mapping to the genome were kept for further analysis. The reads mapped to the *C. carpio* genome were subsequently analysed to annotate rRNA, tRNA, snRNA, snoRNA, and non-coding RNA sequences by blasting against the Rfam (11.0, http://rfam.xfam.org) and GenBank (http://www.blast.nvbi.nlm.nih.gov/) databases. The remaining sequences were identified as the conserved miRNAs in carp by blasting against miRBase 21.0 allowing no more than two mismatches. Existing carp miRNAs referring to *C. carpio* miRNA were included in the miRBase with no base mismatch. The sequences that did not match existing or conserved miRNAs were used to identify potentially novel miRNA candidates (Griffiths-Jones, 2006; Pearson, 1991). Novel miRNA candidates were identified by folding the flanking genome sequence of unique small RNAs using MIREAP (https://sourceforge.net/projects/mireap/). The enrichment level of each miRNA was identified by counting the number of reads in each sample. To identify differentially-expressed miRNAs within the nine libraries, the frequency of miRNA counts was normalized as transcripts per million (TPM). The TPM values were calculated as follows: normalized expression, TPM = (actual miRNA count/number of total clean reads) × 1,000,000. Only the miRNAs with over 2-fold changes in the two compared samples were considered differentially-expressed miRNAs (P < 0.05) (Audic et al., 1997). A positive value represents up-regulation of a miRNA, while a negative value indicates down-regulation.

### Prediction of miRNA Targets

Target genes of miRNAs were predicted using RNAhybrid (v2.1.2) + svm light (v6.01), miRanda (v3.3a) and Targetscan software. The overlap of the predicted results from the three programs was considered to represent the final result of predicted target mRNAs.

### Gene Ontology (GO) and Pathway Analysis of Atrazine-Responsive mRNA Targets

Pathway analysis of the predicted target mRNAs was performed using the Kyoto Encyclopedia of Genes and Genomes (KEGG) pathway database (http://www.genome.jp/kegg/pathway.html) (Kanehisa et al., 2008). To classify the selected genes into groups with similar patterns of expression, each gene was assigned to an appropriate category, according to its main cellular function. To determine the biological phenomena target mRNAs were involved in, the DAVID (http://david.abcc.ncifcrf.gov/home.jsp) functional annotation clustering tool was used.

### QPCR for Validation of miRNAs

The expression profiles of six randomly-selected miRNAs were investigated with qRT-PCR to validate their expression changes. Total RNA (500 ng) was converted to cDNA using miScript reverse transcriptase mix (Qiagen, Valencia, CA, USA) according to the manufacturer’s instructions. QRT-PCR was carried out using an Applied Biosystems 7300 Real-Time PCR System according to the standard protocol. CDNA samples were diluted to 1:150; 5 μL were used for each real-time PCR reaction. The 20-μL PCR mixture included 10 μL SYBR Premix Taq (2X), 0.4 μL miRNA-specific forward primers (10 μM), 0.4 μL miScript universal primer (10 μM), and 1 μL PCR template (cDNA). The PCR thermal program was 50 °C for 2 min, followed by 40 cycles of 95 °C for 2 min, 95 °C for 15 s, and 60 °C for 30 s. Melting curve analysis was performed after amplification. Standard curves for endogenous control and for all miRNAs were constructed using serial dilutions of a pooled cDNA sample. Standard curves were used to determine the quantity of the selected miRNAs and reference genes. Relative miRNA expression levels were calculated using the 2^−ΔΔCt^ method. Each sample was run in triplicate. SnRNA U6 was used as an endogenous control for QPCR of miRNAs.

## RESULTS

### Construction of cDNA Libraries for Sequencing and Small-RNA Discovery

We constructed nine cDNA libraries of small RNAs using pooled total RNAs from gonad tissues exposed to atrazine or control tissues collected from primordial gonad (PG) and from juvenile gonad stage carps. After filtering out low quality sequences, 5**’** and 3**’** adapters, and reads < 18 nt, A total of 10,281,292, 10,086,295, 11,985,647, 10,080,133, 11,724,632, 11, 604, 659, 11, 502, 749, 11, 030, 073, and 11, 282, 882 clean reads were obtained from the nine libraries. Solexa sequencing was then performed for further analysis (Table 1). After comparing the small-RNA sequences with NCBI GenBank and RFam, we removed known types of RNA sequences including rRNA (3.84, 8.80, 8.84, 41.84, 16.31, 5.34, 0.85, 4.42 and 50.12%, respectively), small nuclear RNA (snRNA), small nucleolar RNA (snoRNA), and tRNA (2.16%; 1.84%; 1.08%; 0.47%; 1.08%; 2.72%; 0.58%; 2.79%; and 0.37%), and repeat sequences. Because the genome of common carp is available, the clean reads of small RNAs from the nine libraries were mapped to the common carp genome with miRDeep2 software. A total of 4,895,831 (82.84%); 4,657,428 (84.16%); 7,083,013(84.91%); 5,536,128 (90.12%); 6,104,227 (86.69%); 5,627,337 (82.83%); 5,334,147 (79.12%); 5,337,365 (79.15%) and 7,463,759 (89.37%) of miRNA clean reads were mapped to the common carp genome. The length distribution of the high-quality reads had different trends in the samples within the nine libraries. In the case of PG-CK samples, two peaks of length were observed at 22 nt and 27 nt. However, the size distribution of 21–23 nt increased and the size distribution of 26–29 nt decreased, after exposed to atrazine for 8 h and 24 h (Fig. 1). In the case of IIX-CK samples, higher miRNA mapped rates were observed in small RNAs of 26–28 nt in length. The size distribution of 21–23 nt increased and the size distribution of 26–29 nt decreased after exposure to atrazine for 8 h and 24 h (Fig. 1). In the case of IIC-CK samples, higher miRNA mapped rates were observed in small RNAs of 21–23 nt in length. The size distribution of 21–23 nt decreased and the size distribution of 27–29 nt increased after exposure to atrazine for 8 h and 24 h (Fig. 1). MiRNAs in small RNAs of 26–29 nt in length corresponded to Piwi-interacting RNAs (piRNAs) (Fig. 1), which are endogenous small non-coding RNA molecules 26–31 nt in length. Various studies have shown that Piwi–piRNA complexes are essential in gene silencing and in transposon regulation during germ cell differentiation and gonad development in animals (Klattenhoff and Theurkauf, 2008; Grentzinger et al., 2012; Kawaoka et al., 2012).

**Fig. 1.**
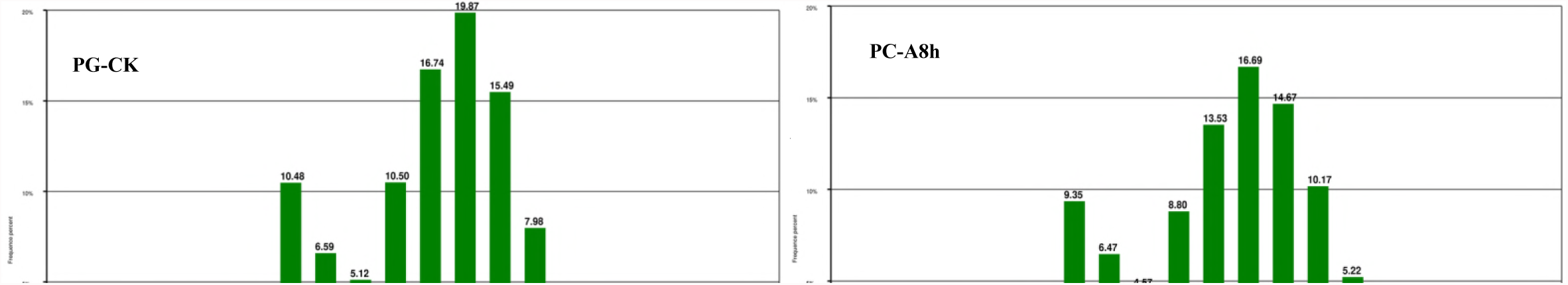
Length distribution of miRNA sequences from Yellow River carp in primordial gonad control (PG-CK), primordial gonad exposed to atrazine for 8 h (PG-A8h), primordial gonad exposed to atrazine for 24 h (PG-A24h), juvenile ovary control (IIC-CK), juvenile ovary exposed to atrazine for 8 h (IIC-A8h), juvenile ovary exposed to atrazine for 24 h (IIC-A24h), juvenile testis control (IIX-CK), juvenile testis exposed to atrazine for 8 h (IIX-A8h), juvenile testis exposed to atrazine for 24 h (IIX-A24h).

**TABLE I.**
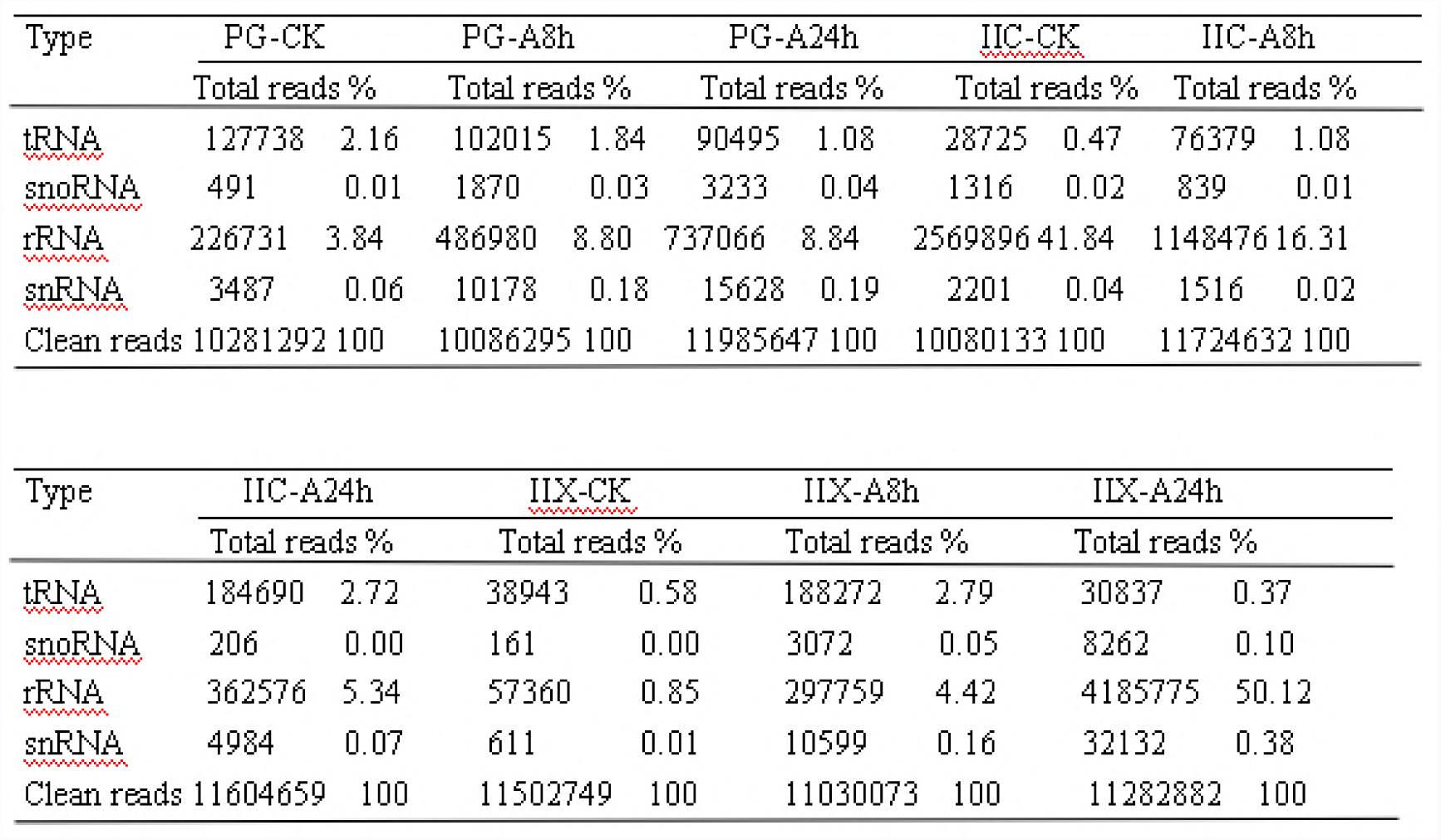
Distribution of sequenced clean reads

### Identification of miRNAs

To identify miRNAs in the gonad of the Yellow River carp exposed or not to atrazine, the clean reads were used and the miRNAs identified by comparison to the deposited miRNAs from miRBase. Mireap_v0.2 software was used for secondary structure prediction of novel miRNA. There was a total of 4,443 miRNAs that were identified, including 3795 existing miRNAs, and 648 conserved miRNAs. Among the existing and conserved miRNAs, 7 miRNAs (ccr-miR-26a, ccr-miR-10b, ccr-miR-143, ccr-miR-181 a, ccr-miR-100, ccr-miR-22a, and ccr-miR-92a) were the most abundant (TPM > 10,000) in all samples (TPM = Readout × 1,000,000 / Mapped reads).

### Validation of miRNAs with qRT-PCR

To validate the results of Solexa sequencing, qRT-PCR was used to test six randomly-selected (ccr-miR-24, ccr-miR-146a, ccr-miR-192, ccr-miR-21, ccr-miR-143, and ccr-miR-454b) miRNAs. According to sequence analysis, from the miRNAs selected for comparison, three miRNAs (ccr-miR-146a, ccr-miR-21, and ccr-miR-454b) were up-regulated in juvenile ovary gonad at 24 h whereas three miRNAs (ccr-miR-24, ccr-miR-192, and ccr-miR-143) were down-regulated in juvenile ovary at 24 h of atrazine exposure. The relative expression levels of all six miRNAs were consistent with the sequencing data (Fig. 2), indicating the reliability of the miRNA expression and correlation analysis based on small-RNA sequencing.

**Fig. 2.**
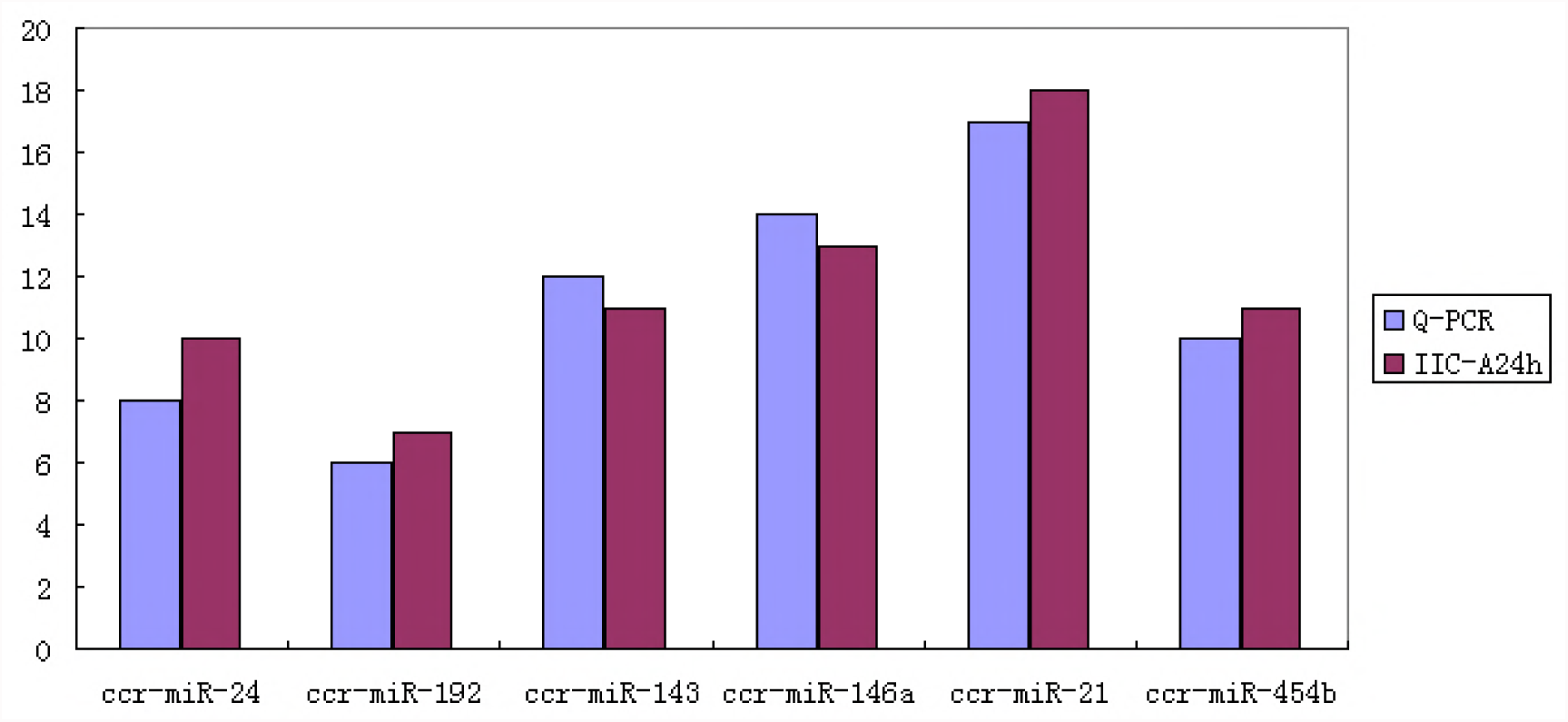
Real-time quantitative PCR gene expression analysis of six randomly-selected miRNAs. Gene expression was normalized to the level of U6 snRNA.

### Effects of Atrazine Exposure on miRNA Expression in PG of Yellow River Carp

Primordial gonad is a crucial stage of sex differentiation, because of the formation of primordial germ cell. A comparative analysis of miRNA expression profiles with or without atrazine exposure may reveal miRNAs with important roles in early gonad differentiation. The results showed that atrazine exposure resulted in the altered expression of a larger number of miRNAs in PG compared with control. Atrazine exposure not only affected the total number of detectable miRNAs, but also the expression levels of miRNAs. After atrazine exposure for 8 h and 24 h, we observed different patterns of differentially-expressed miRNAs in PG of carp. Compared with the control group, 277 miRNAs were up-regulated and 334 miRNAs were down-regulated after atrazine exposure for 8 h. A significant difference in miRNA expression was observed between samples from atrazine exposure for 24 h and unexposed controls, 181 miRNAs were up-regulated and 1,056 miRNAs were down-regulated (Fig. 3). The most significantly down-regulated miRNAs were miR-205, miR-184 and miR-203b-3p, which were down-regulated by 7.15, 3.61 and 3.35 fold, respectively. The most significantly up-regulated miRNAs were miR-7132, miR-135c, and miR-187 which were up-regulated by 8.70, 2.88 and 2.48 fold, respectively (Table 2). Atrazine exposure for 24 h had a greater effect on carp PG miRNA expression than the exposure for 8 h. The number of miRNAs with altered expression after atrazine exposure was higher at 24 h than at 8 h. However, the extent of change varied among the miRNAs. For example, the expression levels of miR-135c and miR-73 8 increased significantly (2.21- and 2.47-fold, respectively), whereas the expression levels of miR-203a decreased significantly (12.0-fold). Similarly, the changes in miRNA expression in PG varied between the unexposed control and atrazine exposure for 8 h or 24 h. For example, miR-135c was up-regulated by 2.2-fold after atrazine exposure for 8 h and was up-regulated by 2.8-fold after atrazine exposure for 24 h. MiR-122 was up-regulated by 1.4-fold after atrazine exposure for 8 h, but was down-regulated by 2.9-fold after atrazine exposure for 24 h. The miRNAs that were significantly altered in PG after exposure to atrazine may thus be involved in sex differentiation and development, and their importance in sex differentiation mechanisms needs to be clarified.

**Fig. 3.**
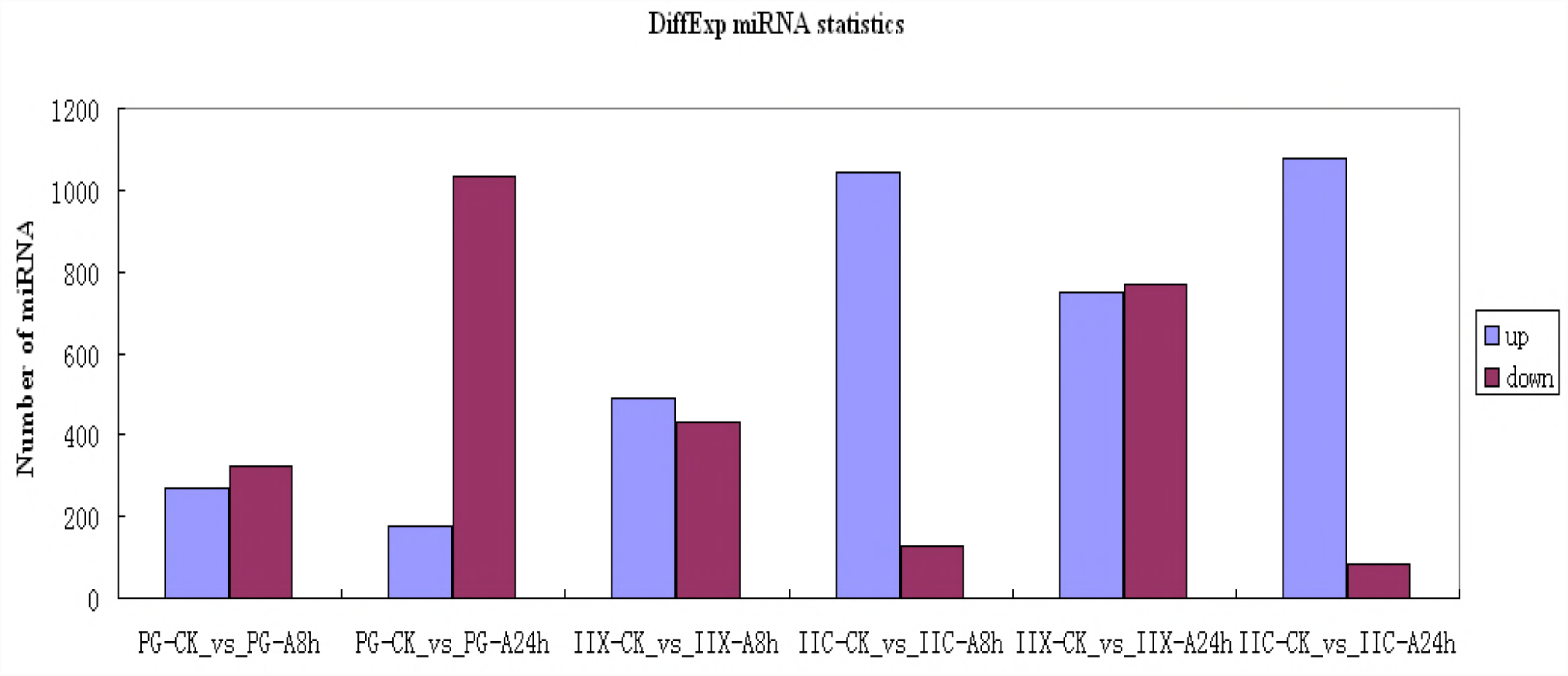
Differential expression of miRNAs in Yellow River carp. Greater than 2-fold change while P < 0.05

**TABLE 2.**
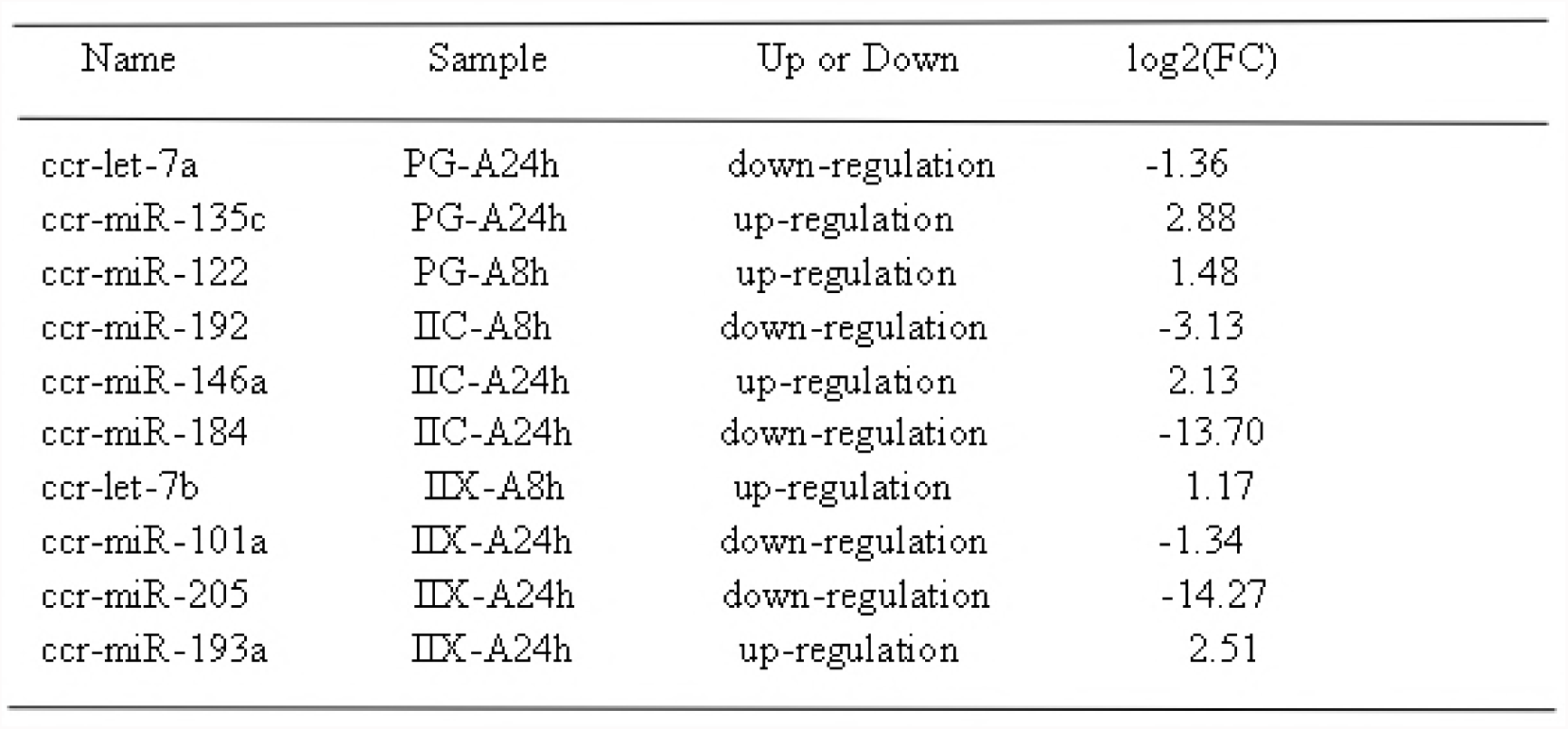
miRNAs with significant expression alterations after atrazine exposure in Yellow River Carp

### Effects of Atrazine Exposure on miRNA Expression in Juvenile Gonad of Yellow River Carp

We observed patterns of differentially-expressed miRNAs in juvenile gonad (stage II ovary and stage II testis) of carp after atrazine exposure for 8 h and 24 h, especially in juvenile ovary (Fig. 3). In juvenile ovary, 1053 miRNAs were up-regulated and 132 miRNAs were down-regulated after atrazine exposure for 8 h, relative to unexposed controls. Relative to the control group, 1085 miRNAs were up-regulated and 84 miRNAs were down-regulated after atrazine exposure for 24 h. The most significantly down-regulated miRNAs were miR-184, miR-214 and miR-122, which were down-regulated by 13.69, 13.21 and 12.40 fold respectively. The most significantly up-regulated miRNAs were miR-17-3p, miR-454a, and miR-454b which were up-regulated by 2.95, 2.49 and 2.42 fold respectively. In juvenile testis, 561 miRNAs were up-regulated and 434 miRNAs were down-regulated after atrazine exposure for 8 h, relative to the control group. Compared with the control group, 775 miRNAs were up-regulated and 799 miRNAs were down-regulated after atrazine exposure for 24 h. The most significantly down-regulated miRNAs were miR-205, miR-194, and miR-122, which were down-regulated by 14.27, 13.59, and 11.81 fold, respectively. The most significantly up-regulated miRNAs were miR-489, miR-738, and miR-193a, which were up-regulated by 10.61, 4.53, and 2.50 fold, respectively. Atrazine exposure for 24 h had a greater effect on juvenile testis miRNA expression than 8 h exposure. In addition, atrazine treatment led to a larger number of miRNAs with altered expression in juvenile testis, than in juvenile ovary. The number of down-regulated miRNAs was higher in juvenile testis than in ovary which is consistent with the feminizing effects of atrazine.

The extent of expression change varied among miRNAs. For example, in juvenile ovary, the miR-301a and miR-17-3p expression levels decreased by 1.38- and 2.95-fold after atrazine exposure for 24 h, respectively. In contrast, the miR-101b expression level decreased by 1.01-fold. In juvenile testis, the miR-193a and miR-146a expression levels increased by 2.50- and 1.71-fold, respectively, after atrazine exposure for with 24 h. In contrast, the miR-122 expression levels decreased by 11.81-fold. Similarly, the changes in miRNA expression of juvenile ovary and testis varied between unexposed controls and atrazine exposure for 8 h or 24 h. For example, ccr-miR-210 was down-regulated by 1.36-fold after atrazine exposure for 8 h, and was down-regulated by 2.06-fold after atrazine exposure for 24 h in juvenile ovary. Ccr-miR-192 was down-regulated by 3.12-fold after atrazine exposure for 8 h, and was down-regulated by 4.07-fold after atrazine exposure for 24 h (Table 2). In juvenile testis, ccr-miR-205 was down-regulated by 3.50-fold after atrazine exposure for 8 h but was down-regulated by 14.27-fold after atrazine exposure for 24 h (Table 2). The miRNAs that were significantly altered in juvenile gonad after exposure to atrazine may thus be involved in sex differentiation and development, and their importance in sex differentiation mechanisms needs to be clarified.

### Expression Patterns of miRNAs at Different Gonad Developmental Stages in Yellow River Carp

Trend analysis of miRNA expression after exposure to atrazine for 8 h and 24 h, at different developmental stages, was conducted. In PG, we identified eight different expression patterns (Fig. 4), including 25 miRNAs that were up-regulated and 214 that were down-regulated during atrazine exposure (Fig. 4, profiles 3, 0). Expression of 232 miRNAs, such as miR-1 and miR-133a-3p, increased after exposure for 8 h, but decreased at 24 h (Fig. 4, profile 5). In contrast, 129 miRNAs, including miR-29a and miR-29b, showed the opposite expression pattern during atrazine exposure (Fig. 4, profile 2). In juvenile ovary, 8 different expression patterns (Fig. 4) were identified, including 440 miRNAs that were up-regulated and 26 that were down-regulated during atrazine exposure (Fig. 4, profiles 7, 0). Expression of 157 miRNAs, such as mir-202-y and mir-27c-5p, increased after exposed for 8 h, but decreased at 24 h (Fig. 4, profile 5). In contrast, 70 miRNAs, including miR-155 and miR-92b, showed the opposite expression pattern during atrazine exposure (Fig. 4, profile 2). In juvenile testis, we also identified eight different expression patterns (Fig. 4), including 68 miRNAs that were up-regulated and 73 that were down-regulated during exposure (Fig. 4, profiles 7, 0). Expression of 117 miRNAs, such as mir-15a and mir-16a, increased after atrazine exposure for 8 h, but decreased at 24 h (Fig. 4, profile 5). In contrast, 41 miRNAs, including miR-144 and miR-148, showed the opposite expression pattern during atrazine exposure (Fig. 4, profile 2).

**Fig. 4.**
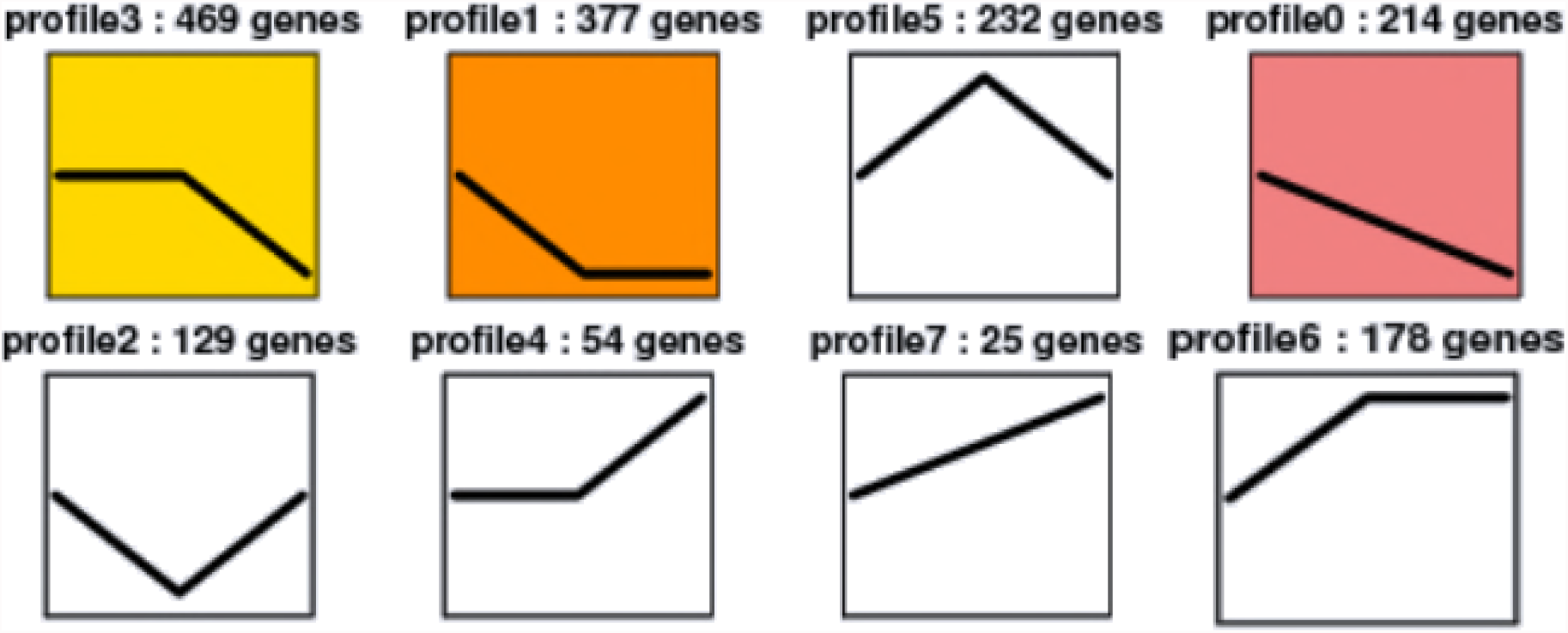
Trend analysis of miRNA expression profiles after exposure to atrazine in Yellow River Carp

In this study, miRNAs targeting male-biased genes showed an upward trend. In PG, miR-499, which was predicted to target *sox9*, increased after exposure to atrazine (Fig. 4, profile 7). *Gsdf* was the predicted target of miR-146a and of miR-22a, which also increased after exposure to atrazine (Fig. 4 profile 7). The expression profiles of miR-72-x and miR-212-y, which were predicted to target *dmrt*, were also consistent with the above miRNAs which predicted male-biased target genes (Fig. 4, profile 7). In juvenile ovary, novel-m3245-5p, which was predicted to target *sox9*, increased after exposure to atrazine (Fig. 4, profile 7). *Gsdf* was the predicted target of novel-m0192-3p and novel-m0514-3p, which also increased after exposure to atrazine (Fig. 4, profile 7). MiR-454a and miR-454b, which were predicted to target *atm*, increased after exposure to atrazine (Fig. 4, profile 7). The expression profiles of novel-m0515-3p and novel-m0080-5p, which were predicted to target *dmrt*, were also consistent with the above miRNAs which predicted male-biased target genes (Fig. 4, profile 18). In juvenile testis, novel-m3312-3p, which is predicted to target *sox9*, increased after exposure to atrazine (Fig. 4, profile 7). *Atm*, which was the predicted target of novel-m0167-3p and novel-m0417-3p, also increased after exposure to atrazine (Fig. 4, profile 7).

In contrast, miRNAs targeting female-biased genes showed a downward trend. Expression levels of novel-m0101-3p and novel-m3450-3p in PG, miR-101b in juvenile ovary, and miR-203b-3p in juvenile testis, all of which were predicted to target *Smad4*, decreased after exposure to atrazine (Fig. 4, profile 0). The most abundant differentially-expressed miRNAs after exposure to atrazine in PG, juvenile ovary and juvenile testis were let-7a, miR-143, and miR-125b, all of which decreased significantly during atrazine exposure.

These results suggest that these miRNAs may influence gonad development.

### Identification and Signalling Analysis of Target Genes of Differentially-Expressed miRNA

To identify potential targets of differentially-expressed miRNAs, involved in sex differentiation and development after atrazine exposure, we performed target-gene prediction based on the common carp (*C. carpio*) genome sequence (http://www.carpbase.org/). A total of 26,299 genes were predicted to be the possible targets of 4353 differentially-expressed miRNAs that were commonly expressed in all atrazine exposure samples. Functional annotation using KEGG identified 239 annotated signalling pathways, including at least 11 pathways involved in reproductive biology: transforming growth factor-β (*TGF-β*) signalling, *Wnt* signalling, oocyte meiosis, mitogen-activated protein kinase (*MAPK*) signalling, Notch signalling, p53 signalling, gonadotropin-releasing hormone (*GnRH*) signalling, RNA polymerase, steroid hormone biosynthesis, estrogen signalling pathway, and metabolism of xenobiotics by cytochrome P450. Interestingly, the target genes of 790 miRNAs belonged to the *MAPK* signalling pathway, which plays an important part in virtually every step of spermatogenesis in the testis. The *MAPK* signalling pathway is also involved in the acrosome reaction in the female reproductive tract before fertilization of the ovum (Huang et al., 2011). *Wnt* signalling is known to be involved in mammalian reproduction (Kobayashi et al., 2011), and in zebrafish sex determination (Chang et al., 2013). We detected 415 miRNA targets belonging to the *Wnt* signalling pathway, 30 belonging to *NF-kappa B* signalling pathway, and 133 belonging to p53 signalling pathway. *Wnt* signalling pathway, *NF-kappa B* signalling pathway and p53 signalling pathways were associated with sex differentiation in zebrafish (Chang et al., 2013). Target genes predicted to belong to the three pathways in our study may be involved in sex differentiation and gonad development in Yellow River carp. Moreover, we identified 245 miRNA targets belonging to the *TGF-β* signalling pathway, and 179 belonging to the Notch signalling pathway. In addition, we also identified 31 miRNA targets belonging to oestrogen signalling pathway, which may play an important role in hormone regulation.

To determine the key biological process of the putative target genes related to atrazine exposure, GO analysis was performed. The identified biological processes that the putative target genes were classified into include reproduction, reproductive process, response to stimulus, developmental process, and growth, which were all mechanisms related to sex differentiation and gonad development. The results showed possible relationships between atrazine, putative targets and gonad development, and suggested that atrazine may have effect on sex differentiation and gonad development.

We analysed the relationships between differentially-expressed miRNAs and their putative target genes. *Foxl2*, *stat1*, *sf1*, *dmrt* and *gsdf* have been shown to be key factors in early ovary differentiation (Ijiri et al., 2008; Nagahama et al., 1997). We also analysed *smad3*, *smad4*, *sox9*, and *atm*, which are also known to be responsible for gonad differentiation. We found that these genes were predicted targets of many miRNAs, which could thus negatively regulate these target genes. Given the important roles of steroid hormones in reproduction and sexual dimorphism in fish, we analysed the relationships between miRNAs, including *hsd11b* and *hsd3b* (which encode key enzymes in the steroid hormone biosynthesis pathway), mRNAs, and the steroid hormone biosynthesis pathway. Atrazine has endocrine-disrupting effects by altering the HPG axis (Trentacoste et al., 2001). We analysed genes that have critical roles in the regulation of the HPG axis including *ER1*, *ER2*, *AR*, and *CYP19A1*.

In the PG, a higher number of miRNAs targeting female-biased genes were down-regulated. MiR-135c which was significantly down-regulated by 2.21-fold and 2.88-fold after exposure to atrazine for 8 h and 24 h, respectively, were predicted to target *ER*, *foxl2*, and *CYP19A*. *Gsdf* was the predicted target of miR-132a, miR-146a, miR-210, and miR-22a which were also down-regulated. Our results indicated that atrazine can promote early gonad-determining genes by down-regulating miRNAs. miR-205, which was predicted to target *atm*, *EGF*, *bcl2*, *BMP1* (bone morphogenetic protein 1), was significantly up-regulated by 7.23-fold for 8 h and 7.15-fold for 24 h, respectively. miR-132a, which targeted *dmrt2*, was up-regulated by 1.22-fold after exposure to atrazine for 24 h, but it was not at 8 h. miR-499, which also targeted *dmrt2*, was up-regulated by 1.46-fold for 8 h and 1.57-fold for 24 h, respectively. After exposure to atrazine for 24 h, miR-202x, and miR-374-y, which were predicted to target *smad3*, were up-regulated and down-regulated, respectively. *Hsd11b* was predicted to target miR-216-x and miR-342-y. *Hsd3b* was the predicted target of let-7-z. *Stat1* was predicted to be the target of miR-13 5c and miR-430, *Sf1* of miR-154-y and miR-395 8-y, and *Sox9* of miR-499. These results illustrate the possible roles of the differentially-expressed miRNAs in PG, after exposure to atrazine during gonad differentiation.

In juvenile ovary, miR-21, which was significantly up-regulated by 2.18-fold after exposure to atrazine for 24 h, was predicted to target *AR* and *atm*. MiR-101 b, which was predicted to target *sf1*, was significantly down-regulated by 1.01-fold. MiR-132a, which was significantly up-regulated by 1.08-fold after exposure to atrazine for 8 h, was predicted to target *AR*, *dmrt2*, *gsdf*, and *atm*. *Smad4* was predicted to target novel-m0048-5p. *Hsd11b* was predicted to target novel-m0305-3p. *Hsd3b* was the predicted target of miR-410-x. *Stat 1* was predicted to be the target of miR-192, *CYP19A* of miR-203a and novel-m0527-3p, and *Sox9* of novel-m0011-5p.

In juvenile testis, miR-181b, and miR-181c, which were significantly up-regulated by 1.04-fold and 1.41-fold, respectively, after exposure to atrazine for 24 h, were predicted to target *dmrt2* and *atm*. miR-146a, which was predicted to target *gsdf* was significantly up-regulated by 1.70-fold. MiR-132a, which was significantly down-regulated by 1.28-fold, was predicted to target *ER*. *Smad4* was predicted to target miR-200b. *Hsd11b* was predicted to target novel-m0305-3p. *Hsd3b* was the predicted target of miR-410-x. *Stat1* was predicted to be the target of miR-192, *CYP19A* of miR-203b-3p and novel-m0693-5p, and *Sox9* of novel-m0081-5p. These results indicate that atrazine promotes the biosynthesis of steroid hormone by altering the miRNAs.

These differentially-expressed miRNAs were also predicted to be involved in many reproductive biology pathways, including steroid metabolic processes, *TGF-β* receptor signalling, *Wnt* signalling, and cell differentiation. Moreover after exposure to atrazine for 24 h, the predicted target genes of the differentially-expressed miRNAs of PG included *cyp51a1*, *hsd3*, *smad4*, *lemd3*, *zranb1*, *tbx6*, *grk6*, *ccna1*, *pcna*, *GATA*, *RBMS1* and *prosapip1*, and some of which, such as *cyp51 a1* and *hsd3*, are gonad development-related genes. Many other miRNAs were also predicted to target genes associated with reproductive processes. miR-205 and miR-135c were predicted to target *bcl2* and *notch2*, which belong to the *TGF-β* signalling and Notch signalling pathways, respectively. miR-205 was predicted to target *pdk1* and *inhibin beta A chain*, which are related to *TGF-β* signalling, and female gonad development, respectively. Although the predicted target genes need to be validated experimentally, these results illustrate some of the possible roles of the differentially-expressed miRNAs in gonad reproductive processes.

## DISCUSSION

MiRNAs are involved in diverse biogenesis pathways and have versatile regulatory functions in differentiation, proliferation, and apoptosis (Bartel 2009). To date only a limited number of studies have investigated miRNA expression alterations in response to exposure to endocrine-disrupting chemical in fish and humans (Avissar-Whiting et al., 2010; Hsu et al., 2009; Jenny et al., 2012; Tilghman et al., 2012; Veiga-Lopez et al., 2013). There are only few reports on the miRNA profiling of fish, in response to atrazine exposure, and no reports in common carp. The investigation into the adverse effects of atrazine exposure on miRNAs is important to reveal the molecular mechanism of gonad differentiation. In the present study, we assessed the potential effects of atrazine on miRNAs in the reproductive system at two developmental stages (PG and II-stage gonad) of Yellow River carp. Primordial germ cell formation is a crucial stage of gonad differentiation, and II-stage gonad is the stage of evident sex differentiation. Comparative analysis of miRNA expression profiles at these two important stages, after exposure to atrazine, is helpful to identify miRNAs that play important roles in gonad differentiation.

In this study, atrazine exposure resulted in significant expression alterations of various miRNAs. Atrazine exposure for 24 h caused more alterations in the expression of miRNAs than exposure for 8 h. Atrazine exposure for 24 h caused more alteration in miRNA expression in juvenile testis than in juvenile ovary. It is thus clear that acute and short-time exposure to atrazine during development can produce adverse effects, as has been suggested before (Kathryn et al., 2016).

Several studies in amphibians have suggested that atrazine is associated with feminization of males in the wild (Hayes et al., 2002; Hayes et al., 2002; Murphy et al., 2006). In field studies, atrazine has repeatedly been associated with the presence of feminized secondary sex characteristics in male frogs (McCoy et al., 2008). In fish, atrazine causes degeneration of interstitial tissue in the testes (Spano et al., 2004) and feminizes the gonads of developing male teleost fish (Tillitt et al., 2008). In addition, embryonic atrazine exposure alters the expression of zebrafish and human miRNAs known to play a role in angiogenesis, cancer, neuronal development, differentiation, and maturation (Sara et.al 2016). In our study, atrazine exposure altered the expression of carp miRNAs that play a role in gonad differentiation and gonad development. A number of miRNAs (including miR-122, let-7, miR-192, miR-21, miR-499, miR-146, miR-101, miR-128, and miR-124) that are highly expressed in adult bighead carp and silver carp were significantly altered in our study (Chi et al., 2011).

Our results suggest that miR-21, let-7, miR-430, miR-181a, and miR-143 may play important roles in gonad differentiation and development in Yellow River carp.

Several studies suggested that miR-21 may play an important role in gonad development. A study reported that in cattle miR-21 was significantly up-regulated in the ovary (relative to testis) suggesting that miR-21 may play a regulatory role in female physiology (McBride, 2012). A previous study indicated that miR-21 plays a role in preventing apoptosis in periovulatory granulosa cells, as they transit into luteal cells (Christenson et al., 2010). Has-miR-21 was also up-regulated by ovarian steroids in mouse granulosa cells and human endometrial stromal cells, and in glandular epithelial cells (Fiedler et al., 2008; Pan et al., 2007). In this study, atrazine exposure did not change the expression of miR-21 in the PG after atrazine exposure, but induced its up-regulation in juvenile ovary and down-regulation in juvenile testis.

The predicted target genes of miR-21 included genes of the *MAPK*, B-cell receptor, *TGF-β*, and apoptotic pathways. This observation suggests that miR-21 may play crucial roles in ovary development, gonad differentiation (Gangaraju and Lin et al., 2009), and endocrine regulation (Eshel et al., 2014; Huang et al., 2011). The predicted target genes of miR-21 in our study were *AR* and *atm*.

Let-7 was another family of miRNAs with altered expression by atrazine exposure. The let-7 family was first discovered and characterized in *Caenorhabditis elegans*, and plays an important role in regulating late developmental events by down-regulating lin-41, and possibly other genes (Pasquinelli et al., 2000). Let-7 was significantly up-regulated after atrazine exposure in the PG and juvenile testis. The predicted target genes of let-7 in our study were *sox9* and *atm*.

The miR-430 family is known to be involved in embryonic morphogenesis and clearance of maternal mRNAs; it is and highly expressed during early zebrafish development (Choi et al., 2007; Giraldez et al., 2005; Giraldez et al., 2006; Inui et al., 2010). MiR-430 has been shown to target chemokine signalling to ensure accurate migration of primordial germ cells (Staton et al., 2011). In our study and miR-430 was down-regulated in PG but not in juvenile gonad, which indicates that miR-430 has an important role in early gonad differentiation of Yellow River carp.

Several reports showed that miR-143 is highly expressed in the juvenile ovary; it is a dominant miRNA in ovaries in cattle, pigs, and yellow catfish (Li et al., 2009; Lau et al., 2014). In this study, miR-143 was highly expressed in juvenile ovary, which is in keeping with previous reports.

The miR-181a family is abundantly expressed in the gonads of tilapia (Hossain et al., 2012), mice (Saunders et al., 2010), and humans (Sirotkin et al., 2009). It was down-regulated in juvenile ovary in the present study. Overall, above results suggest that miR-21, let-7, miR-430, miR-181a, and miR-143 may play important roles in goand differentiation and development in Yellow River carp.

Differentially expressed miRNAs showed a variety of expression patterns at different development stages. Among the 8 different expression patterns, two patterns are particularly worthy of attention, involving miRNAs with expression levels that either increased or decreased significantly after atrazine exposure. MiRNAs whose expression either increased or decreased significantly after atrazine exposure may be direct regulators of gonad differentiation. Samples with the highest number of miRNAs with altered expression were the PG and juvenile ovary exposed to atrazine for 8 h or 24 h. The number of decreased miRNAs was 1,056 in PG, including miRNAs which targets were female-biased. Because miRNAs are negatively correlated with its target genes, this observation suggests that atrazine promotes the expression of female-biased genes by decreasing specific miRNAs in PG, which would result in the differentiation of the gonad to the female phenotype. The juvenile ovaries exposed to atrazine had the highest number of up-regulated miRNAs, including miRNAs whose targets are male-biased. It is thus possible that atrazine represses the expression of male-biased genes by increasing specific miRNAs in juvenile ovary.

The juvenile testis exposed to atrazine had the highest number of miRNAs with altered expression, indicating that this tissue was more sensitive to atrazine, possibly leading to the feminization of males. This observation suggests that these miRNAs may have an important function in the timing of gonad differentiation and development.

Target-gene prediction showed that many of the genes that we identified as targets of the miRNAs that we studied were involved in sex differentiation. Among these predicted genes, *sox9*, *dmrt*, and *gsdf* have been identified as sex-determining genes in fish (Diego et al., 2015; Myosho et al., 2012). For example, *Hsd11b* and *hsd3b* encode key enzymes in the steroid hormone biosynthesis pathway. These genes may participate in steroid hormone synthesis, gonad function, and mechanisms of sex differentiation, and may play a vital role in developmental timing. However, further studies are needed to confirm the interactions and functions of miRNA and target genes. In addition, the results also show that atrazine has oestrogenic effects down-regulating male-biased genes (such as *dmrt* and *atm*) through specific miRNAs up-regulation, and up-regulating female-biased genes (such as *foxl2*) through specific miRNAs down-regulation.

Previous studies showed that atrazine exposure can significantly reduce synthesis, secretion, and the circulating levels of androgens in fish (Moore et al., 1998; Spano et al., 2004), amphibians (Hayes et al., 2002; Hayes et al., 2010), reptiles (Rey et al., 2009), and mammals (Friedmann et al., 2002; Stoker et al., 2000), and also in birds to a lower extent (Wilhelms et al., 2006). The endocrine-disrupting effects of atrazine are primarily due to alterations of the HPG axis (Cooper et al., 2000; Foradori et al., 2009, 2013; Weber et al., 2013; Wirbisky et al., 2016a). However, atrazine’s mechanism of action is not well-understood, it has been proposed that atrazine up-regulate aromatase expression (Caron-Beaudoin et al., 2016; Sanderson et al., 2000, 2001, 2002). Aromatase up-regulation leads to increased conversion of androgens into oestrogens (Laville et al., 2006). In the present study, we analysed genes that regulate hormone biosynthesis in the HPG axis, including *ER1, ER2*, *AR*, and *CYP19A1*. MiR-122, which targets *ER1* and *ER2*, was down-regulated by atrazine. MiR-21, which targets *AR* was up-regulated in PG by atrazine. MiR-203a, which targets *CYP19A1*, was down-regulated in PG by atrazine. Our results indicate that atrazine can up-regulate aromatase expression through specific miRNAs, which is consistent with previous studies.

We tested the hypothesis that atrazine has endocrine-disrupting effects by altering genes of the HPG axis through its corresponding miRNAs. In the PG, atrazine affects sex differentiation mainly through altering upstream genes involved in gonad differentiation. In juvenile ovary or testis, atrazine affects the gonad development mainly through altering hormone generation and the expression of hormone receptor genes. Further studies are needed to investigate the mechanisms and roles of miRNAs in the regulation of genes during gonad differentiation and development.

In summary, atrazine exposure caused significant alterations in miRNAs expression at the crucial stages of carp gonad development. Target genes of differentially-expressed miRNAs are key factors in early ovary differentiation or play an important role in the formation of germ cells. In addition, our results indicate that atrazine up-regulates aromatase expression through specific miRNAs, supporting the hypothesis that atrazine has endocrine-disrupting effects, altering the expression of genes of the HPG axis through its corresponding miRNAs.

## COMPETING INTEREST

**The authors have declared that no competing interests exist.**

Luo, X. Y., Sunohara, Y., Matsumoto, H., 2004. Fluazifop-butyl causes membrane peroxidation in the herbicide-susceptible broad leaf weed bristly starbur (Acanthospermum hispidum). Pestic. Biochem. Physiol. 78 (2), 93 – 102.

